# Large X-linked palindromes undergo arm-to-arm gene conversion across Mus lineages

**DOI:** 10.1101/800185

**Authors:** Callie M. Swanepoel, Emma R. Gerlinger, Jacob L. Mueller

**Affiliations:** Department of Human Genetics, University of Michigan Medical School, Ann Arbor, MI USA

## Abstract

Large (>10kb), nearly-identical (>99% nucleotide identity), palindromic sequences are enriched on mammalian sex chromosomes. Primate Y-palindromes undergo high rates of arm-to-arm gene conversion, a proposed mechanism for maintaining their sequence integrity in the absence of X-Y recombination. It is unclear whether X-palindromes, which can freely recombine in females, undergo arm-to-arm gene conversion and, if so, at what rate. We generated high-quality sequence assemblies of *Mus molossinus* and *Mus spretus* X-palindromic regions and compared them to orthologous *Mus musculus* X-palindromes. Our evolutionary sequence comparisons found evidence of X-palindrome arm-to-arm gene conversion at rates comparable to rates of autosomal allelic gene conversion in mice. Mus X-palindrome genes also exhibit higher than expected sequence diversification, indicating gene conversion may facilitate the rapid evolution of palindrome-associated genes. We conclude that in addition to maintaining genes’ sequence integrity via sequence homogenization, arm-to-arm gene conversion can also rapidly drive genetic evolution via sequence diversification.

## Main Text

Approximately a quarter of the human Y chromosome is comprised of large (>10kb), nearly-identical (>99%) palindromic sequences (Skaletsky, et al. 2003). The presence of Y-palindromes in multiple primate species reflects duplication events that occurred prior to their speciation. Despite the agedness of Y-palindromes, comparative sequencing of primate Y-palindromes revealed lower levels of sequence divergence between paralogous palindrome arms (within a species) than between orthologous arms (between species), suggesting that ongoing arm-to-arm gene conversion homogenizes the sequence within in paralogous palindrome arms (Rozen, et al. 2003). Primate Y-palindrome arm-to-arm gene conversion is estimated to be 3X higher than allelic gene conversion (Rozen, et al. 2003; Halldorsson, et al. 2016). Since most of the Y chromosome does not recombine with the X chromosome, Y-palindrome arm-to-arm gene conversion has been proposed to maintain the sequence integrity of the testis-specific genes harbored within Y-palindromes (Rozen, et al. 2003; Skaletsky, et al. 2003).

Large palindromes are also overrepresented on the human and mouse X chromosomes (Warburton, et al. 2004; Mueller, et al. 2008; Mueller, et al. 2013). Given that the X chromosome freely recombines in females, it is unknown whether X-palindromes also undergo arm-to-arm gene conversion and, if so, at what rate. We conducted a comparative sequence analysis of X-palindromic regions across three mouse lineages, *Mus musculus*, *Mus molossinus* and *Mus spretus*, to assess whether X-palindromes undergo frequent arm-to-arm gene conversion.

Due to the complexity of large, nearly identical palindromes, producing high-quality assemblies is imperative to confirm their palindromic orientation and confidently determine sequence divergence. We investigated singleton X-palindromes, as opposed to arrays of palindromes, because they are commonly found on mammalian sex chromosomes and we can more accurately assess their rates of gene conversion (Skaletsky, et al. 2003; Warburton, et al. 2004; Mueller, et al. 2013). Since whole-genome shotgun approaches are unable to assemble large segmental duplications with high levels of nucleotide identities, a large-insert clone-based sequencing strategy is necessary for accurate assembly of large palindromes (Treangen and Salzberg 2011; Hughes and Rozen 2012; Vollger, et al. 2019). Thus, we generated high-quality X-palindrome assemblies from a single haplotype by sequencing *M. molossinus* and *M. spretus* male (XY) bacterial artificial chromosome (BAC) clones with PacBio long read sequencing. Twelve *M. molossinus* BACs and three *M. spretus* BACs were sequenced, spanning six palindromes (300kb of palindromic sequence) in *M. molossinus*, and three palindromes (136kb of palindromic sequence) in *M. spretus* (Supplementary Table 1). Additional X-palindromes may exist in *M. molossinus* and *M. spretus*, but were not detectable in our screens (see Materials and Methods). Each sequenced BAC spanned either part or the entirety of the palindrome (arm sizes ranging from 10-65kb) and also included single copy sequence flanking the palindromes (Supplementary Table 1, Supplementary Fig. 1). Based on 1.0% sequence divergence between paralogous arms and a 5.4 × 10^−9^ substitution rate per nucleotide per generation in mice, *M. musculus* X-palindromes could have arisen via recent segmental duplications ~178,000 years ago (Uchimura, et al. 2015) (Fig. 1). However, the presence of orthologous palindromes across all three Mus lineages, having diverged up to 3 million years ago (MYA), indicates X-palindromes are not the result of recent segmental duplications (Fig. 1) (Lundrigan and Tucker 1994; Harr, et al. 2016).

**Fig 1.**
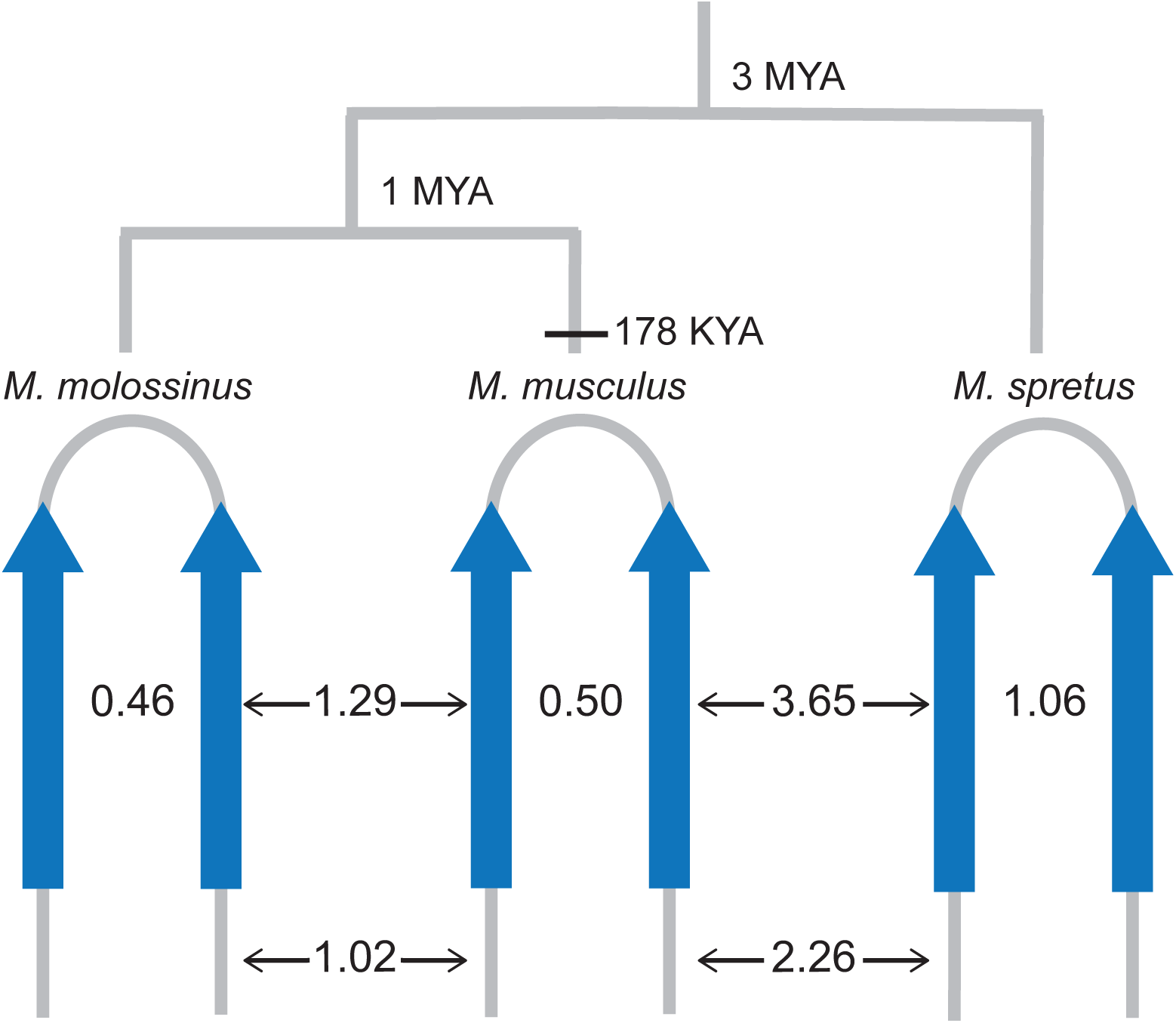
Mus X-palindromes, conserved across three Mus lineages, exhibit paralogous versus orthologous sequence divergence indicative of ongoing palindrome arm-to-arm gene conversion. A phylogenetic tree representing the evolutionary relationship between *M. musculus*, *M. molossinus* and *M. spretus* (Lilue, et al. 2018). X-palindromes are present in all three lineages, indicating their sequence identity is not a consequence of recent segmental duplication ~178,000 years ago (YA). Schematics of palindromes in each lineage represent the total percent divergence between paralogous (within a lineage) and orthologous (across lineages) palindrome arms for all X-palindromes analyzed. The total percent divergence between single copy flanking sequence is also shown.

If X-palindromes are not the result of recent segmental duplications, then palindrome arm-to-arm gene conversion could explain the higher than expected sequence identity between paralogs. Palindrome arm-to-arm gene conversion could occur intrachromosomally in the male germline and both intra- and inter-chromosomally in the female germline. To detect X-palindrome arm-to-arm gene conversion, we compared paralogous and orthologous palindrome arm sequence divergence across all three Mus lineages (Fig. 1, Supplementary Tables 2, and 3). Ongoing gene conversion is characterized by low levels of sequence divergence between paralogous palindrome arms relative to between orthologous palindrome arms. We found this pattern of sequence divergence to be true across all three Mus lineages (p-value = 2.2×10^−16^, exact binomial test), consistent with ongoing arm-to-arm gene conversion (Fig. 1, Table 1). The high identity between X-palindrome arms could alternatively be explained by their presence in regions of the X chromosome with low mutation rates. In this case, we would also expect a similar low mutation rate in sequences flanking the palindromes. We found that the total sequence divergence of orthologous X-palindrome flanking sequence is 1.02% and 2.26% for *M. molossinus* and *M. spretus*, respectively (Fig. 1, Table 1, Supplementary Table 3), which is in agreement with estimates of X chromosome divergence in *M. molossinus* (0.72% div.) and *M. spretus* (1.5% div.) (Abe, et al. 2004; Lilue, et al. 2018). From these data, we conclude that X-palindromes are not in regions of low mutation rates and are undergoing arm-to-arm gene conversion across all three Mus lineages.

**Table 1.** Total sequence divergence between paralogous and orthologous palindrome arms

| Comparison | Length (kb) | Total Div. (%) | 95% CI |
| --- | --- | --- | --- |
| *M. musculus* (arm-to-arm) | 205 | 0.50 | 0.47-0.53 |
| *M. molossinus* (arm-to-arm) | 154 | 0.46 | 0.42-0.49 |
| *M. spretus* (arm-to-arm) | 66 | 1.06 | 0.98-1.14 |
| *M. musculus* vs *M. molossinus* |  |  |  |
| Palindrome arm | 270 | 1.29 | 1.25-1.33 |
| Single-copy sequence | 76 | 1.02 | 0.95-1.08 |
| *M. musculus* vs *M. spretus* |  |  |  |
| Palindrome arm | 132 | 3.65 | 3.55-3.76 |
| Single-copy sequence | 42 | 2.26 | 2.12-2.41 |

We next sought to determine the X-palindrome arm-to-arm gene conversion rate. Sequence identity between palindrome arms is maintained by a steady-state balance of new variants being introduced and subsequently erased or fixed by arm-to-arm gene conversion (Rozen, et al. 2003). To estimate the rate of X-palindrome arm-to-arm gene conversion, we assume *c* = 2μ/*d* (c is the arm-to-arm gene conversion rate, d is the divergence between arms and μ is the mutation rate at 5.4 × 10^−9^ per nucleotide per generation in mice) (Rozen, et al. 2003; Uchimura, et al. 2015). We find the rate of mouse X-palindrome arm-to-arm gene conversion is: *c* = 2 × 5.4 × 10^−9^/5.0 × 10^−3^ = 2.17 × 10^−6^ gene conversions per duplicated nucleotide, per generation. The X-palindrome gene conversion rate is thus similar to the estimated allelic gene conversion rate of 3.04 × 10^−6^ per nucleotide per generation, based upon recent high-resolution mapping studies in mice (Li et al. 2018). Our findings suggest X-palindrome arms behave more similar to homologous chromosomes than paralogous duplications, which tend to have lower gene conversion rates (Harpak, et al. 2017).

Y-palindrome arm-to-arm gene conversion is thought to protect palindrome-associated genes by replacing deleterious mutations. If this is the case, then one would expect arm-to-arm gene conversion events to be biased towards ancestral variants. This is observed in primate Y-palindromes, where divergence between orthologous palindrome arms is significantly lower than divergence of single-copy sequence on the Y chromosome (Rozen et al. 2003; Hallast et al. 2013). In contrast, we find significantly greater divergence between orthologous X-palindrome arms relative to that of single-copy X-linked sequence across all three Mus lineages (p = 2.2×10^−16^, exact binomial test; Table 1, Supplementary Table 3). This pattern suggests there is selection for new variants (derived), not ancestral, in X-palindromes, a pattern also observed for a single Y-palindrome in European rabbits (Geraldes et al. 2010).

We reasoned that if there is selection for increased divergence in X-palindromes, then X-palindrome associated genes could show signatures of rapid evolution. We thus examined the sequence evolution of X-palindrome-associated genes by comparing the ratio of nonsynonymous to synonymous substitution rates (K_a_/K_s_) between *Mus musculus* and *Rattus norvegicus* genes in X-palindromes (Table 2), to non-palindromic genes on the X chromosome and across the genome. We found X-palindrome genes have a median K_a_/K_s_ ratio of 0.54 (n=5), which is significantly higher than the median of other X-linked genes (K_a_/K_s_ =0.22 (n=741)) (p = 0.0287) and genome-wide (K_a_/K_s_ =0.11 (n=11,503)) (Gibbs, et al. 2004). Our findings suggest that the presence of two palindromic gene copies and arm-to-arm gene conversion is an important mechanism to promote the rapid evolution of their associated genes.

**Table 2.** Sequence evolution of X-palindrome associated genes between *Mus musculus* and *Rattus norvegicus*

| X-palindrome Gene | A | S | K<sub>a</sub> | K<sub>s</sub> | K<sub>a</sub>/K<sub>s</sub> |
| --- | --- | --- | --- | --- | --- |
| *4930567H17Rik* | 359.2 | 159.8 | 0.49 | 1.10 | 0.44 |
| *Mageb5* | 834.9 | 260.1 | 0.10 | 0.17 | 0.60 |
| *Zxdb* | 2051.6 | 546.4 | 0.62 | 1.78 | 0.35 |
| *Gm773* | 479.7 | 138.3 | 0.09 | 0.16 | 0.54 |
| *Xlr5a* | 533.2 | 171.8 | 0.13 | 0.25 | 0.54 |

A = mean number of nonsynonymous sites
S = mean number of synonymous sites
K<sub>a</sub>= mean number of nonsynonymous substitutions per nonsynonymous site
K<sub>s</sub>= mean number of synonymous substitutions per nonsynonymous site

We conclude that X-palindromes undergo arm-to-arm gene conversion at rates concordant with allelic gene conversion, potentially facilitating the rapid evolution of their associated genes. Why are the arm-to-arm gene conversion rates of primate Y-palindromes higher than the rates of primate allelic and mouse X-palindrome gene conversion? One possibility is that there is stronger selection for palindrome arm-to-arm gene conversion on the Y chromosome, than on the X chromosome, since X chromosomes can undergo allelic gene conversion in females. A second possibility is that long palindrome arm lengths (up to 1.5Mb) on the primate Y, compared to the much shorter mouse X-palindromes (10-65kb), may more readily facilitate gene conversion (Skaletsky, et al. 2003). A third possibility is that there are inherent differences in the location and rate of gene conversion between primates and mice, warranting assessment of mouse Y-palindrome and human X-palindrome arm-to-arm gene conversion rates. To distinguish between these possibilities and build on the exclusively evolutionary-based gene conversion estimates of palindromes to date, a chromosome-wide assessment of X- and Y-palindrome gene conversion rates in modern day humans and mice is necessary.

## Materials and Methods

### Identification of Mus molossinus and Mus spretus BACs

*M. musculus* contains eight singleton X-palindromes with arms spanning 13-65kb in length (Kruger, et al. 2018). We screened *M. spretus* (CHORI-35 SPRET/Ei) male mouse (XY) high-density filters of the BAC library using hybridization probes. Hybridization probes were designed within palindrome arms and spacers of the eight *M. musculus* X-palindromes (Supplementary Table 4). In cases where the *M. musculus* and *M. spretus* hybridization probe sequences differ, we converted probe sequence nucleotide variants between *M. musculus* and *M. spretus* to *M. spretus* sequence. 66 BACs were positive from filter hybridization screening, 34 of which were positive upon PCR screening using primers within the spacers or flanking the palindrome arms (Supplementary Table 5). Of the 34 BACs, three spanned palindrome arms and were selected for sequencing (Supplementary Table 1).

To identify *M. molossinus* BACs, we screened BAC-end sequences from a male BAC library (Abe, et al. 2004). To identify *M. molossinus* BACs that span the eight X-palindromes arms, we used position and orientation of *M. molossinus* BAC-end sequences that mapped to unique positions on the X chromosome. We identified 17 BACs spanning the eight mouse X-palindromes, 12 of which were selected for sequencing (Supplementary Table 1).

### BAC sequencing and assembly

BAC sequencing and assembly was performed as previously described (Vollger, et al. 2019). Briefly, BAC DNA from clones was isolated using a High Pure Plasmid Isolation Kit from Roche Applied Science (PN: 11754785001) following the manufacturer’s instructions, using 6 ml of LB media with chloramphenicol as a selective marker. Non-overlapping BACs were pooled at equal molar amounts before library preparation. Approximately 1 μg of DNA per BAC was pooled and sheared using a Covaris g-TUBE. Libraries were processed using the PacBio SMRTbell Template Prep kit following the protocol “Procedure and Checklist—20 kb Template Preparation Using BluePippin Size-Selection System” with the addition of barcoded SMRTbell adaptors. Library size was measured using a FEMTO Pulse, and libraries were pooled at equal molar amounts before size selection. The pooled library was size-selected on a Sage PippinHT with a start value of 12,000 and an end value of 50,000. BACs were sequenced on a PacBio Sequel with version 3.0 chemistry on one SMRT cell and then demultiplexed using LIMA in SMRTlink6.0. Demultiplexed reads were run through the CCS algorithm in SMRTlink6.0. CCS reads were filtered for contaminating *E. coli* reads, and the resulting filtered fasta was used as input for assembly using Canu v1.8. Canu contigs were then split on the vector sequence, had the vector removed, and were stitched back together with Sequencher. These contigs were then polished with Arrow, and visualized with Parasight (http://eichlerlab.gs.washington.edu/jeff/parasight/index.html) to check for even coverage (Supplementary Figure 2). All BACs were checked against their end sequences and/or the mouse genome assembly on the UCSC genome browser to confirm BAC identity. To assemble overlapping BACs into larger contigs, BACs were aligned using default parameters in Sequencher. Altogether, 1.76Mb and 0.53Mb of sequence was assembled in *M. molossinus* and *M. spretus*, respectively.

### Identification of palindromes and their boundaries in sequenced BACs

To identify palindromes within *M. molossinus* or *M. spretus* BACs or overlapping BACs (contigs), a pair-wise comparison of each contig against itself was performed using MacVector and displayed graphically as a dot-plot and as an alignment with the coordinates of palindrome arm boundaries. Palindromic sequence was defined as a region of alignment greater than 2kb in length.

### Estimating sequence divergence between orthologous and paralogous palindrome arms

Orthologous and paralogous palindromic sequences were aligned in MEGA using CLUSTAL W with default parameters (Thompson, et al. 1994; Kumar, et al. 2016). *M. molossinus* and *M. spretus* sequence was aligned to the *M. musculus* (mm10) reference genome using BLAT (Kent 2002) to identify orthologous palindromic and single-copy flanking sequence. For palindrome arms consisting of multiple fragments, separated by single-copy sequence (Supplementary Figure 1), each fragment was individually aligned to its respective paralogous or orthologous sequence. To estimate divergence, p-distance, or the proportion of nucleotide sites at which two sequences are different (variant sites) was calculated using default parameters in MEGA (Supplementary Tables 2 and 3) (Kumar, et al. 2016). The number of variant sites in each alignment was then calculated by multiplying the p-distance by the total number of aligned sites (Supplementary Tables 2 and 3). We estimated 95% confidence intervals and compared the divergence of paralogs vs orthologs and palindromes vs single copy sequence using a two-sided exact binomial test under the null hypothesis that the proportion of variant sites is the same between the groups compared.

### Estimating K_a_/K_s_ of mouse-rat orthologs

Rat palindrome gene orthologs were identified using the NCBI Refseq gene track from the rn6 reference genome. Rat palindrome genes that were not annotated were identified based on synteny with the mouse reference genome and the presence of predicted rat mRNA orthologous to mouse mRNA. Nucleic acid sequences from mouse and rat were translated to peptide sequences using EMBOSS Transeq and aligned using CLUSTAL Omega (Madeira, et al. 2019). K_a_/K_s_ of mouse-rat palindrome gene orthologs were calculated using the PAL2NAL program with default parameters (Suyama, et al. 2006). Mouse-rat X-linked orthologs, K_a_ and K_s_ values were obtained from the Ensembl database using biomaRt. K_a_/K_s_ of palindrome gene orthologs was compared to the K_a_/K_s_ of X-linked gene orthologs using the Mann-Whitney U test.

## Data Availability

Complete BAC sequences generated in this study are available from NCBI Genbank under accession numbers: MN249934-249948.

## Acknowledgements

We thank Pieter de Joung and BACPAC for screening CH35 BAC libraries. We thank the University of Washington PacBio Sequencing Center: Katherine M. Munson, Alexandra P. Lewis and Melanie Sorensen for generating the PacBio libraries, sequencing and assembly. CMS was supported by the following NIH training grant: “Michigan Predoctoral Training in Genetics (T32GM007544). This work was supported by the National Institutes of Health (grant number HD094736).

