## Supplementary Fig. and Supplementary Tables for "Large X-linked palindromes undergo arm-to-arm gene conversion across Mus lineages"

**Letter**

**Discoveries**

Affiliations:

**This PDF file includes:**

- **Supplementary Tables 1-5**
- **Figs. S1, S2**

Supplementary Table 1. *M. molossinus and M. spretus* sequenced BACs containing X-palindromes

| BAC ID | Mouse Lineage | Region | Assembly length (bp) |
|---|---|---|---|
| MSMg01-501C17 | *M. molossinus* | *Gm773* | 185,880 |
| MSMg01-476N20 | *M. molossinus* | *4930567H17Rik* | 205,955 |
| MSMg01-519D22 | *M. molossinus* | *4930567H17Rik* | 164,511 |
| MSMg01-512B09 | *M. molossinus* | *Xlr5a* | 181,943 |
| MSMg01-46A24 | *M. molossinus* | *Mageb5* | 168,673 |
| MSMg01-120K10 | *M. molossinus* | *Mageb5* | 96,195 |
| MSMg01-294L01 | *M. molossinus* | *Mageb5* | 106,397 |
| MSMg01-546J18 | *M. molossinus* | *Zxdb* | 82,744 |
| MSMg01-128K01 | *M. molossinus* | *Zxdb* | 134,477 |
| MSMg01-355O22 | *M. molossinus* | *Zxdb* | 157,090 |
| MSMg01-150D05 | *M. molossinus* | *3010001F23Rik* | 132,621 |
| MSMg01-444A04 | *M. molossinus* | *3010001F23Rik* | 140,587 |
| CH35-410H24 | *M. spretus* | *4930567H17Rik* | 197,660 |
| CH35-113M24 | *M. spretus* | *Zxdb* | 163,382 |
| CH35-317J20 | *M. spretus* | *Xlr5a* | 172,730 |

Supplementary Table 2. Sequence divergence of paralogous palindrome arms

| Comparison | Palindrome | Arm Length (kb) | Aligned Sites | Variant Sites | Total Div. (%) |
|---|---|---|---|---|---|
| ***M. musculus vs M. musculus*** | *Xlr5a* | 25.6 | 25,530 | 54 | 0.21 |
| | *4930567H17Rik* | 64.5 | 64,120 | 415 | 0.65 |
| | *Zxdb* | 48.6 | 48,494 | 215 | 0.44 |
| | *Gm773* | 10.6 | 10,595 | 34 | 0.32 |
| | *3010001F23Rik* | 28.2 | 28,149 | 117 | 0.42 |
| | *Mageb5* | 28.4 | 28,302 | 185 | 0.65 |
| | **Total** | **205.9** | **205,190** | **1,020** | **0.50** |
| ***M. molossinus vs M. molossinus*** | *Xlr5a* | 26.7 | 26,712 | 7 | 0.03 |
| | *4930567H17Rik* | 30.8 | 30,066 | 252 | 0.84 |
| | *Zxdb** | 20.1 | 20,958 | 93 | 0.45 |
| | *Gm773* | 10.7 | 10,698 | 8 | 0.07 |
| | *3010001F23Rik* | 28.5 | 28,417 | 201 | 0.71 |
| | *Mageb5** | 36.7 | 36,682 | 139 | 0.38 |
| | **Total** | **153.5** | **153,533** | **701** | **0.46** |
| ***M spretus vs M. spretus*** | *Xlr5a* | 26.5 | 25,098 | 278 | 1.11 |
| | *4930567H17Rik* | 25.9 | 25,149 | 391 | 1.55 |
| | *Zxdb** | 15.8 | 15,759 | 30 | 0.19 |
| | **Total** | **68.2** | **66,006** | **699** | **1.06** |

* In cases where we only have partial palindrome sequence for the non-*M. musculus* lineage

Supplementary Table 3. Sequence divergence of orthologous palindrome arms and single-copy flanking sequence

| Comparison | Palindrome | Aligned Sites | Variant Sites | Total Div. (%) |
|---|---|---|---|---|
| ***M. musculus* vs *M. molossinus* (palindrome arm)** | *Xlr5a* | 51,674 | 575 | 1.11 |
| | *4930567H17Rik* | 57,510 | 889 | 1.55 |
| | *Zxdb* | 41,667 | 566 | 1.36 |
| | *Gm773* | 21,388 | 153 | 0.71 |
| | *3010001F23Rik* | 56,785 | 895 | 1.58 |
| | *Mageb5* | 41,047 | 413 | 1.01 |
| | **Total** | **270,071** | **3,491** | **1.29** |
| ***M. musculus* vs *M. molossinus* (single-copy flanking sequence)** | *Xlr5a* | 19,728 | 298 | 1.51 |
| | *4930567H17Rik* | 16,041 | 138 | 0.86 |
| | *Zxdb** | 9,985 | 75 | 0.76 |
| | *Gm773* | 13,978 | 140 | 1.00 |
| | *3010001F23Rik* | 9,884 | 88 | 0.89 |
| | *Mageb5** | 6,435 | 33 | 0.51 |
| | **Total** | **76,051** | **772** | **1.02** |
| ***M. musculus* vs *M. spretus* (palindrome arm)** | *Xlr5a* | 50,084 | 2,284 | 4.56 |
| | *4930567H17Rik* | 50,230 | 1,812 | 3.61 |
| | *Zxdb** | 31,313 | 713 | 2.28 |
| | **Total** | **131,627** | **4,809** | **3.65** |
| ***M. musculus* vs *M. spretus* (single-copy flanking sequence)** | *Xlr5a* | 18,787 | 472 | 2.51 |
| | *4930567H17Rik* | 17,009 | 335 | 1.97 |
| | *Zxdb** | 6,542 | 149 | 2.28 |
| | **Total** | **42,338** | **957** | **2.26** |

* In cases where we only have partial palindrome sequence for the non-*M. musculus* lineage

Supplementary Table 4. Hybridization probes for CHORI-35 SPRET/Ei BAC library screen

| **Hybridization Probe** | **Sequence** |
|---|---|
| *4930567H17Rik*_arm | CTCGTATAATGACACAGAATATCCTTCAGCTACTATTGATAT |
| *4930567H17Rik*_spacer | GCTCTGGTTCACTTAGTAGAAATGATAATATCCTGAGTACAG |
| *Mageb5*_arm | AGATCTTATTGAACCAAAGTACCGGTATAATTTTAGTGATAT |
| *Mageb5*_spacer | TAGCATAAATCTTTAGAGATACTCCAGATTTTAGTTAGGTAC |
| *301000F23Rik*_arm | ATACAAACAGATAAAAACTCTTTGGAATGTATTAACAACACC |
| *301000F23Rik*_spacer | ATAGAAGAAAGATTTTCTGGGATAGATAAGTAAAGAAAGAGA |
| *Xlr5a*_arm | AAGAGGAGACAAAGTGAGGTTTACCATGGAAGTTCTTTTTCC |
| *Xlr5a*_spacer | GTGAATTTGTACATTCTGATGAAGTACATTCTATTGTGAACA |
| *Zxdb*_arm | TGGCATCTTGACTATTGATGTGGCCTCTGTGAACTCGAGCCT |
| *Zxdb*_spacer | AAAAGCAAAATTTTTGTCTAGATTTATCTGCACAGCCAAGAT |
| *Gm773*_arm | CAGTTTCACTGTAGTTTATTATCAGTAATCAGTAACATTTAA |
| *Gm773*_spacer | AGCATAAATTAATAATCCTCTAGCACATTAACCATCTATAGC |

Supplementary Table 5. Primers for PCR screening CHORI-35 SPRET/Ei BAC clones

| Primer | Sequence |
|---|---|
| *Gm773*_5'F | GCTCATCAATGGGAAGGAGA |
| *Gm773*_5'R | TGGACACTTCCAAACCTTCC |
| *Gm773*_spacerF | TCTTGCAAAGGCTCAAACTG |
| *Gm773*_spacerR | TTGCTTTGACTCTGCCACTG |
| *Gm773*_3'F | TGAAGCTCTTTGGAGGCATT |
| *Gm773*_3'R | CCATGCCCTCTGTGGACTAT |
| *3010001F23Rik*_5'F | CAGATCAGAAGGAGCCCAAG |
| *3010001F23Rik*_5'R | CTGTCCATGCCTAATGCAGA |
| *3010001F23Rik*_spacerF | CTAAGCCAGGCCAACTTGAG |
| *3010001F23Rik*_spacerR | GGCCTTCTACACGCAGAAAC |
| *3010001F23Rik*_3'F | CAAAGCTGATCCCCTGTGAC |
| *3010001F23Rik*_3'R | ACTGTGTGCAGCAGATGAGG |
| *4930567H17Rik*_5'F | TGCTGTTTTCTGGATTGCACA |
| *4930567H17Rik*_5'R | CAGTTTTGGTTGCTTGAGGGA |
| *4930567H17Rik*_spacerF | ACGGCTCTGGTTCACTTAGT |
| *4930567H17Rik*_spacerR | TAGGAATCACTTGCATGTCG |
| *4930567H17Rik*_3'F | TTTGGCCTTACCACCTGAAC |
| *4930567H17Rik*_3'R | ATTTCATGGCATGGATTGGT |
| *Xlr5a*_5'F | CATTGAGCAGCAGGAAGGAC |
| *Xlr5a*_5'R | CAGGCATTCACACACTGACA |
| *Xlr5a*_spacerF | TGGAGGCCACTTTGTTGAAT |
| *Xlr5a*_spacerR | GCTGAGTGGCTAATGATACTG |
| *Xlr5a*_3'F | GAAGTGAGGGGTACGATGCT |
| *Xlr5a*_3'R | CGATAGCGTGCCAAGAAGTC |
| *Mageb5*_5'F | AAAGACGCCCAGAGAAGAGT |
| *Mageb5*_3'R | CTCCAAGCGCACAGAGATTC |
| *Mageb5*_spacerF | GTTTGCAGAGTTGTGGACTGATAC |
| *Mageb5*_spacerR | ATCATTTTCCTTGTAGAGAACACAGC |
| *Mageb5*_3'F | TAAATTACTAGTCTCTTCTCAAGATGTCC |
| *Mageb5*_3'R | ACACAATGAGAACTACCAGAGTAGAGG |
| *Zxdb*_5'F | TGATCCAGAGCAGACACCAG |
| *Zxdb*_5'R | TGCCTCGTTACCATTGAACA |
| *Zxdb*_spacerF | CCTATGCATTCCAGTCCTTGT |
| *Zxdb*_spacerR | TTGCGAGTATCTGTGGGTGT |
| *Zxdb*_3'F | AATGGGGAAGGAAAGGAGCA |
| *Zxdb*_3'R | CCTACCACCTGCGGCTAACA |

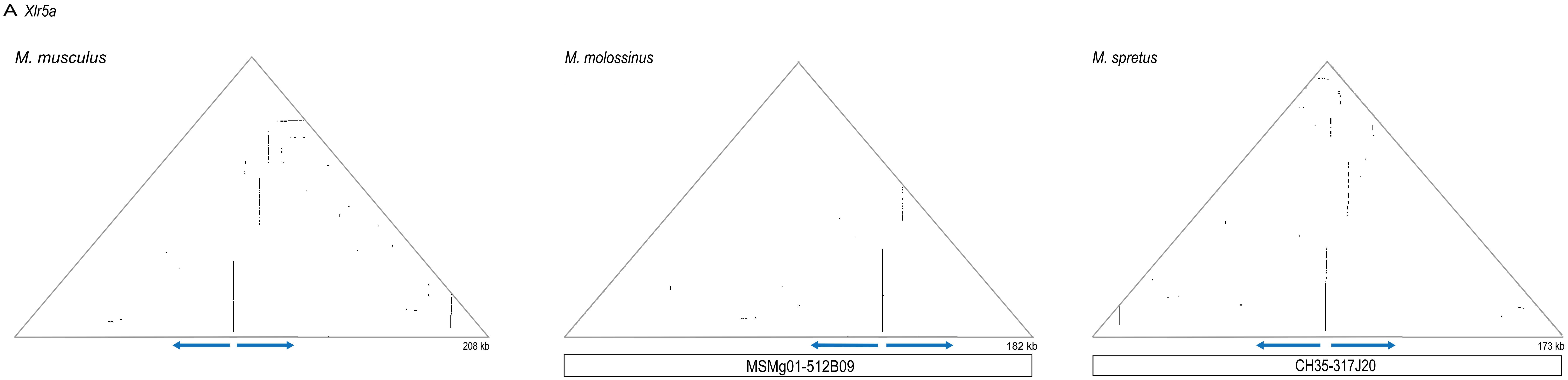
A Xlr5a
M. musculus
208 kb
M. molossinus
182 kb
MSMg01-512B09
M. spretus
173 kb
CH35-317J20

B *4930567H17Rik*

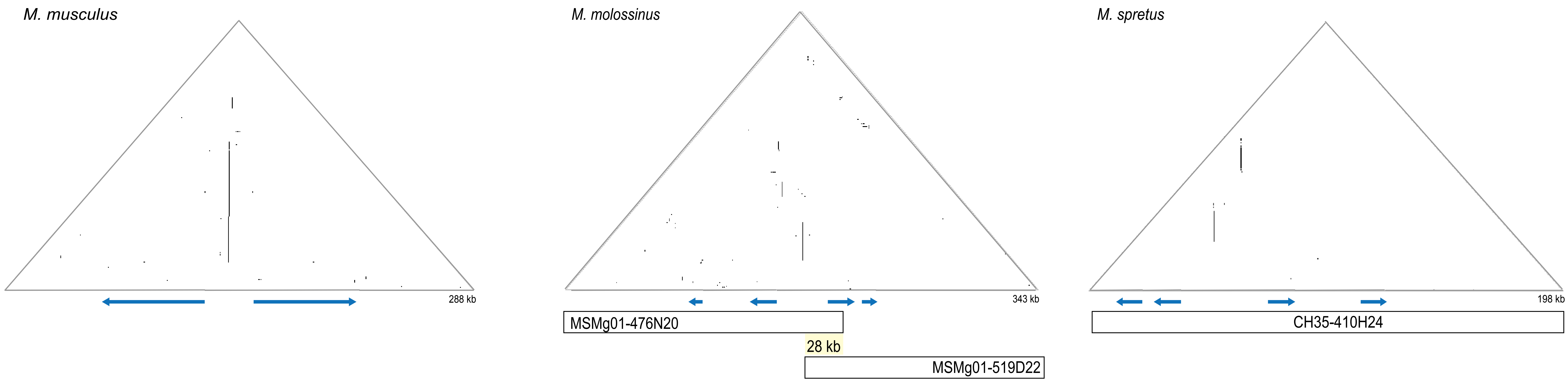

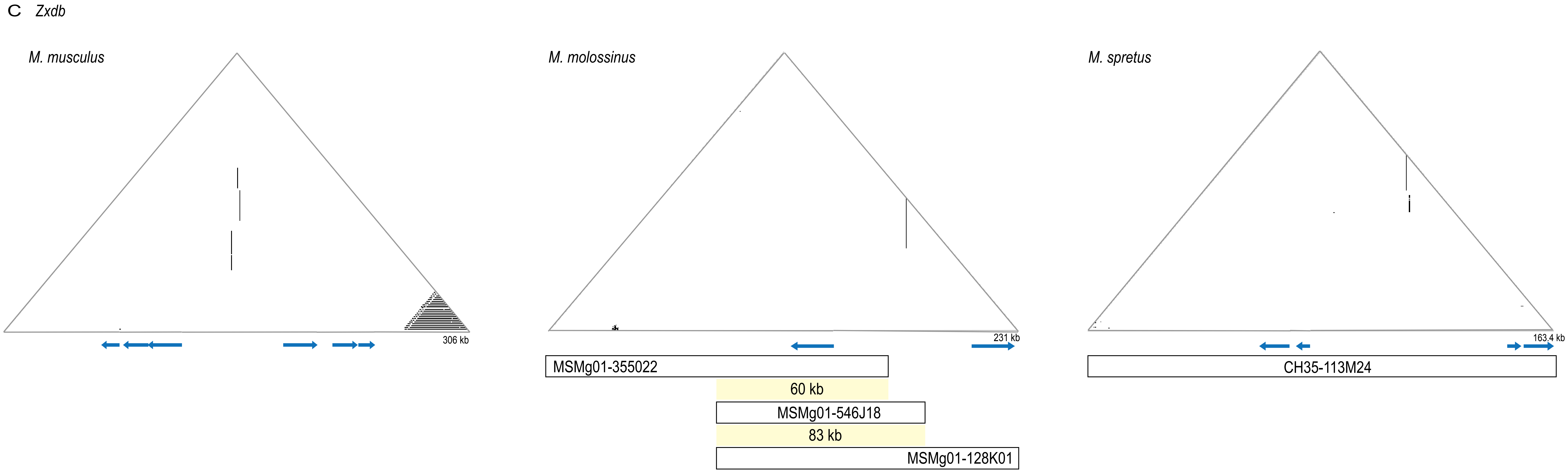
C Zxdb
M. musculus
306 kb
M. molossinus
231 kb
MSMg01-355022
60 kb
MSMg01-546J18
83 kb
MSMg01-128K01
M. spretus
163.4 kb
CH35-113M24

D *Gm773*

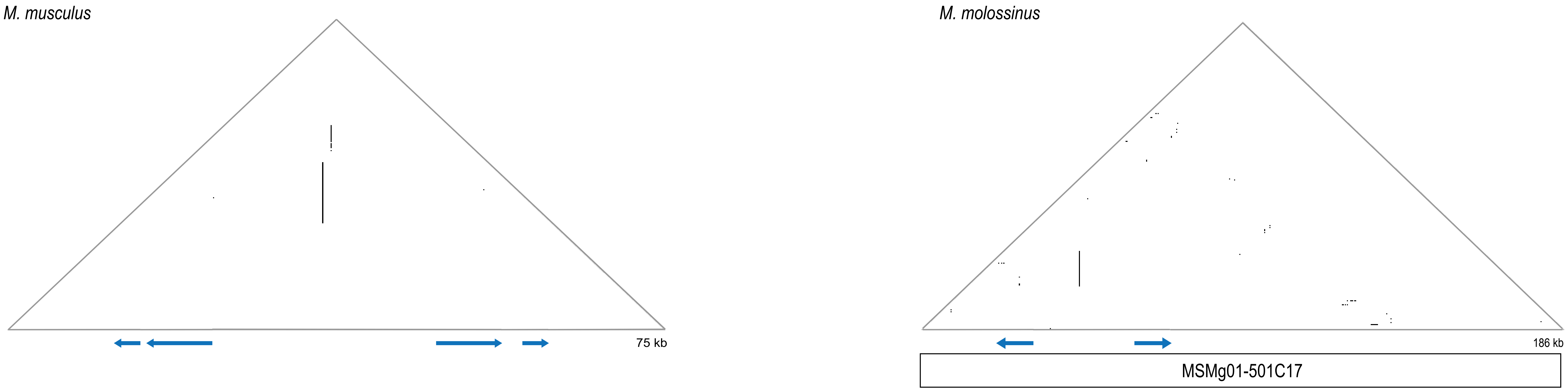

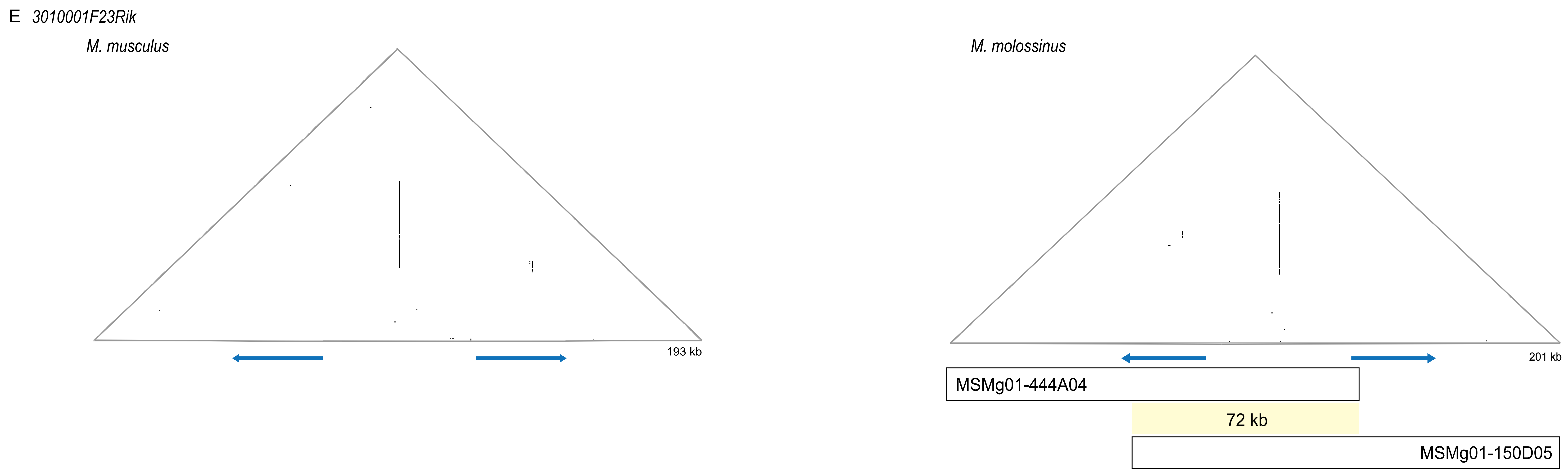
E 3010001F23Rik
M. musculus
193 kb
M. molossinus
201 kb
MSMg01-444A04
72 kb
MSMg01-150D05

F *Mageb5*

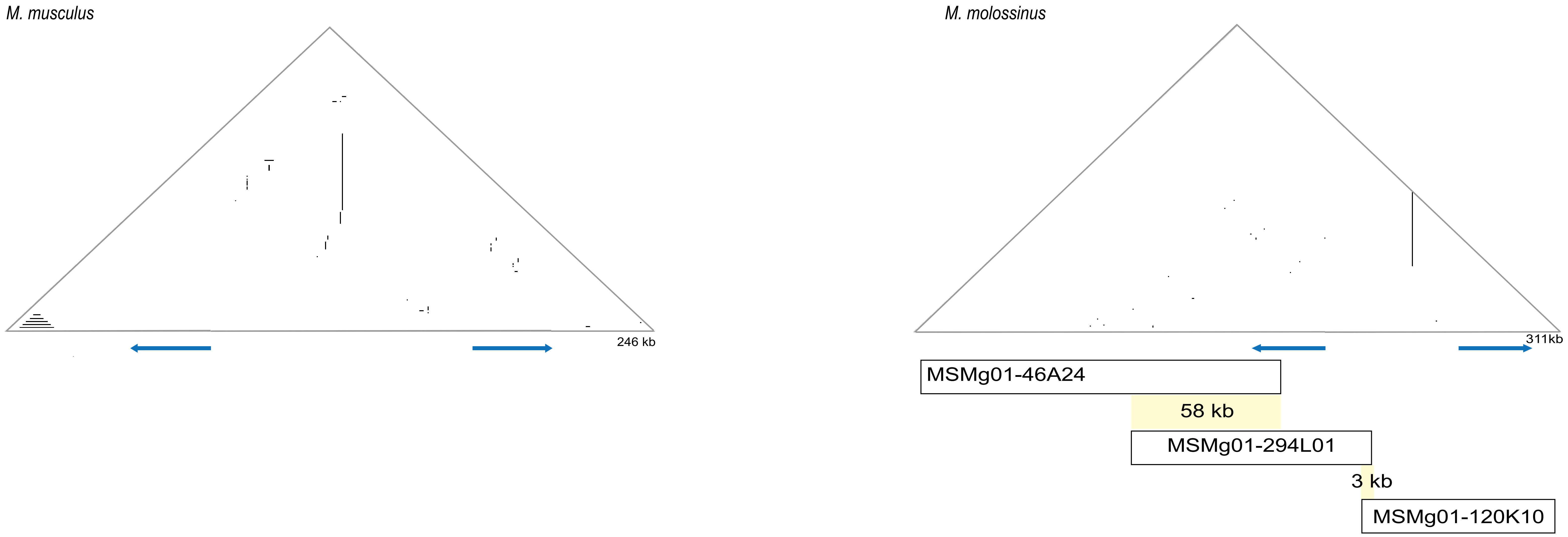
M. musculus
246 kb
M. molossinus
311kb
MSMg01-46A24
58 kb
MSMg01-294L01
3 kb
MSMg01-120K10

**Fig. S1. Self-symmetry triangular dot plots of X-chromosome palindromes in three Mus lineages**. *M. musculus* DNA from the mm10 reference genome or sequenced BACs from *M. Molossinus* and *M. spretus* spanning X-palindromes are plotted against themselves with a sliding window of 100 nucleotides and step size of 1 nucleotide. A 100 nucleotide window that is identical to the sequence it is compared to is represented as a dot and vertical lines represent duplications present in palindromic orientation. Blue arrows below the x-axis represent palindrome arms. Six different palindromic regions were analyzed: A) *Xlr5a*, B) *4930567H17Rik*, C) *Zxdb* gene, D) *Gm773,* E) *3010001F23Rik,* and F) *Mageb5*.

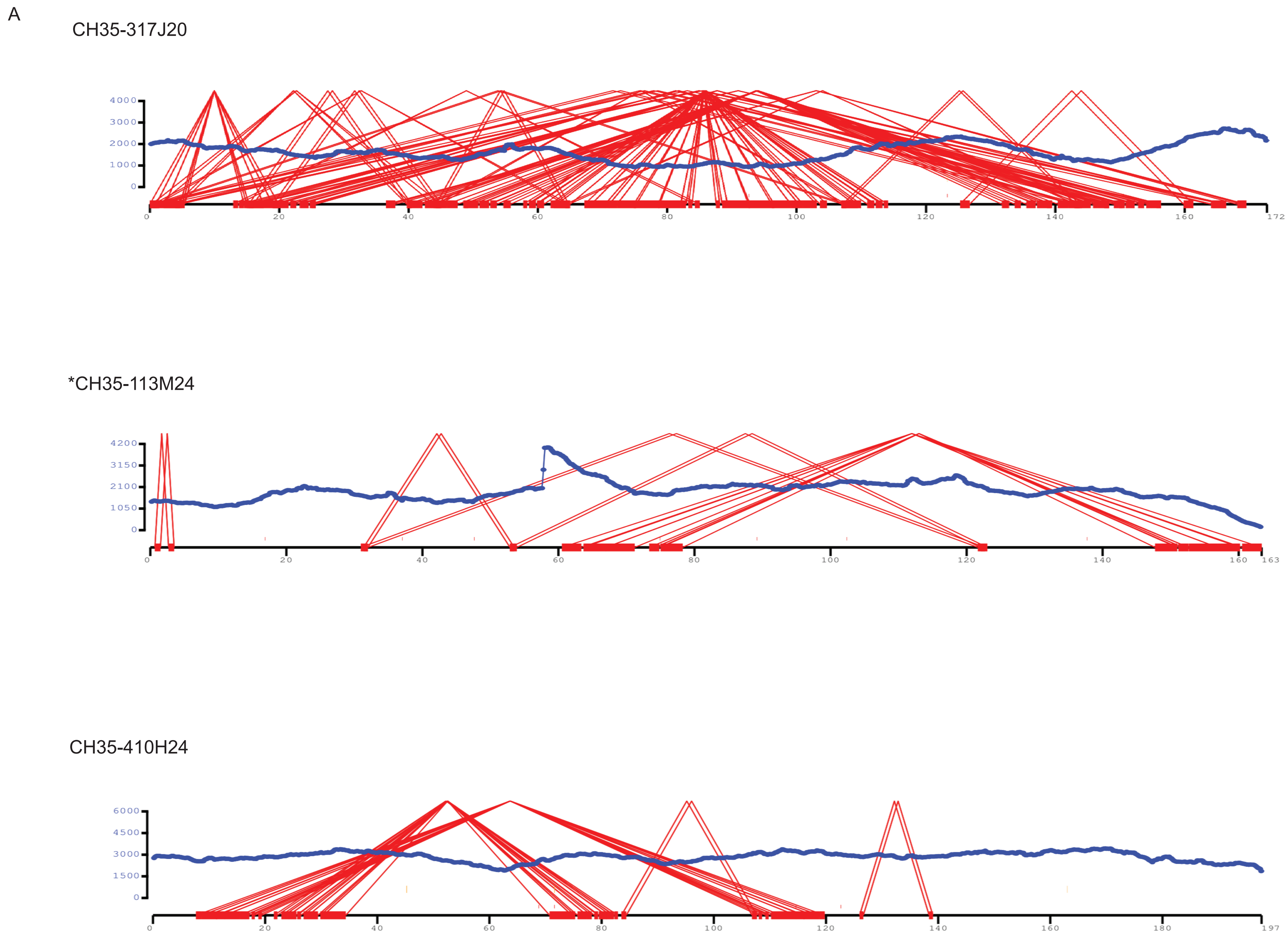
A
CH35-317J20
4000
3000
2000
1000
0
0
20
40
60
80
100
120
140
160
172
*CH35-113M24
4200
3150
2100
1050
0
0
20
40
60
80
100
120
140
160
163
CH35-410H24
6000
4500
3000
1500
0
0
20
40
60
80
100
120
140
160
180
197

B

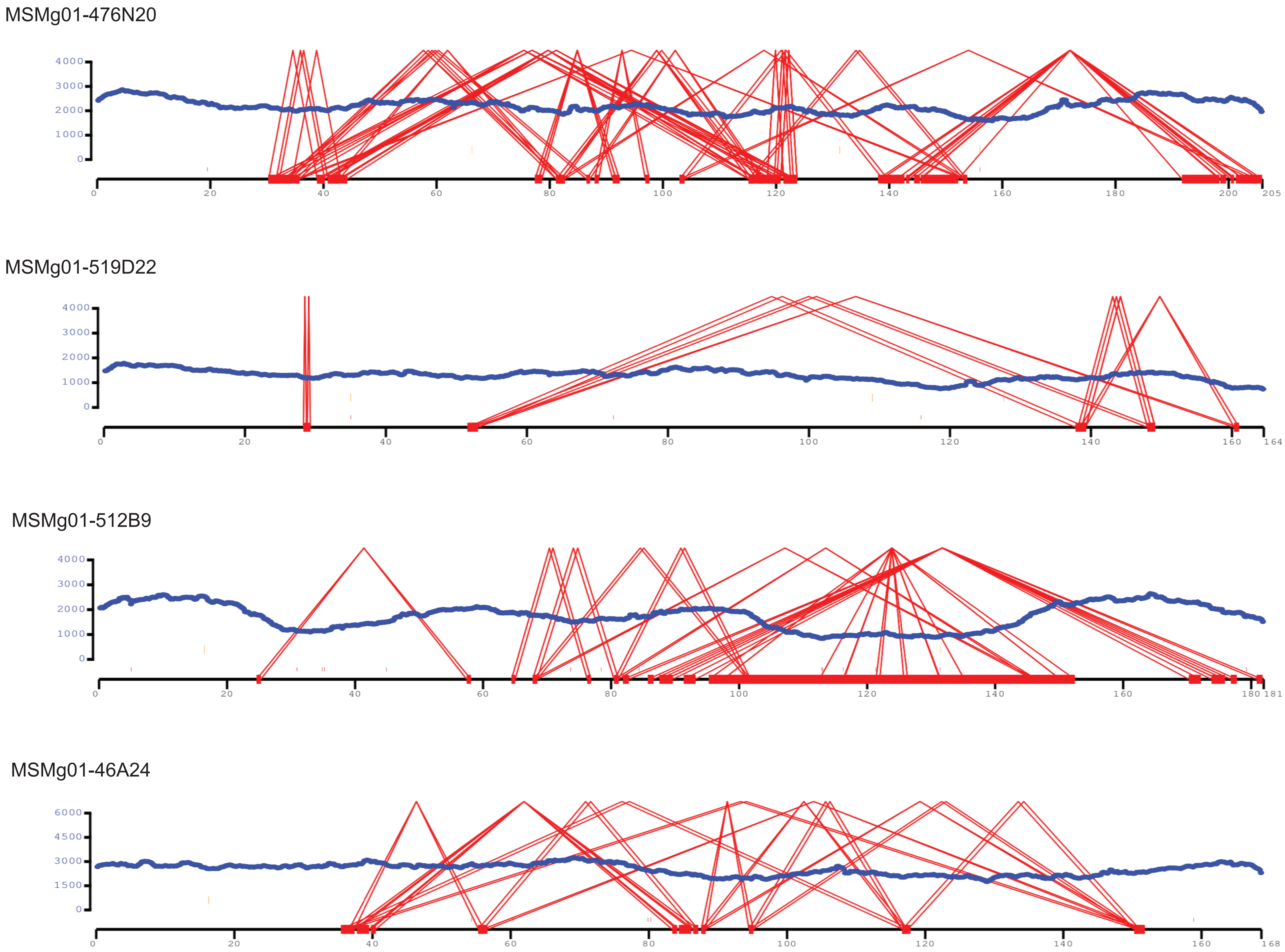

MSMg01-120K10

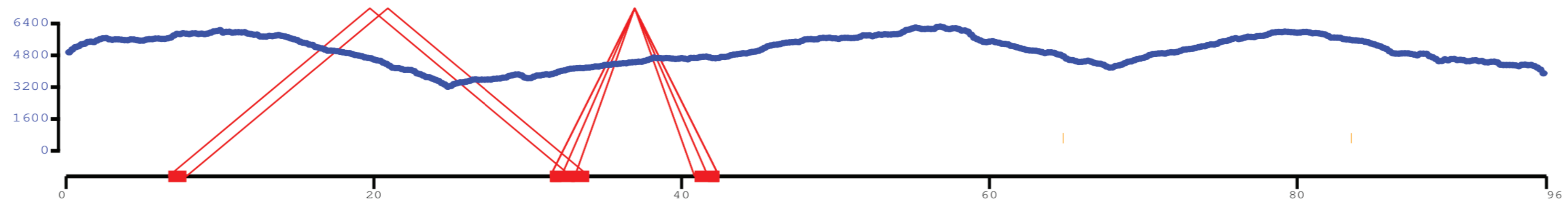


MSMg01-294L01

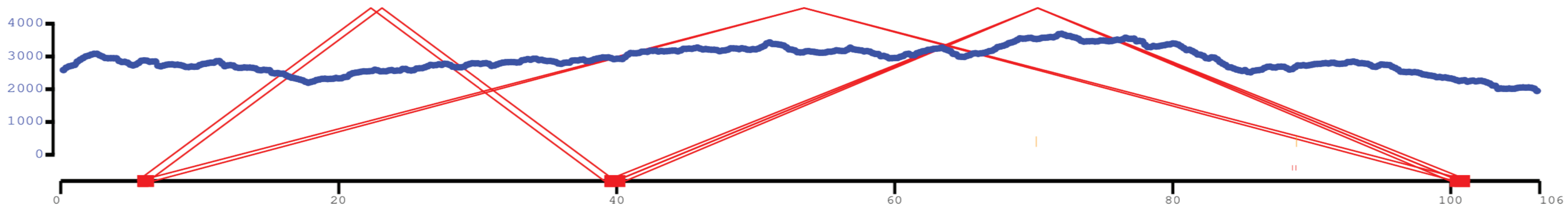


MSMg01-546J18

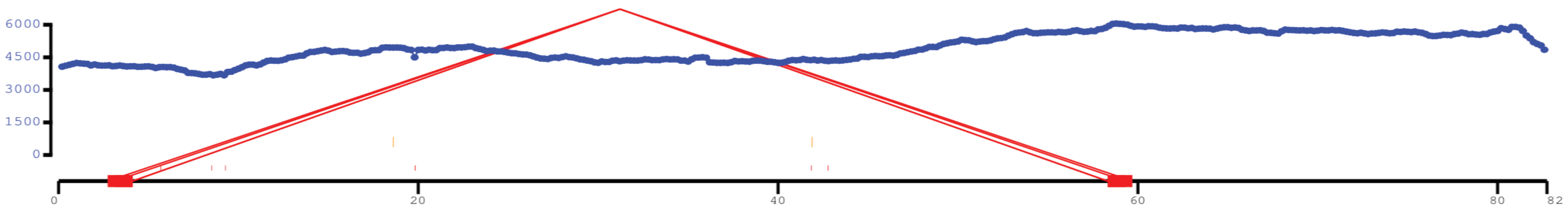


MSMg01-128K01

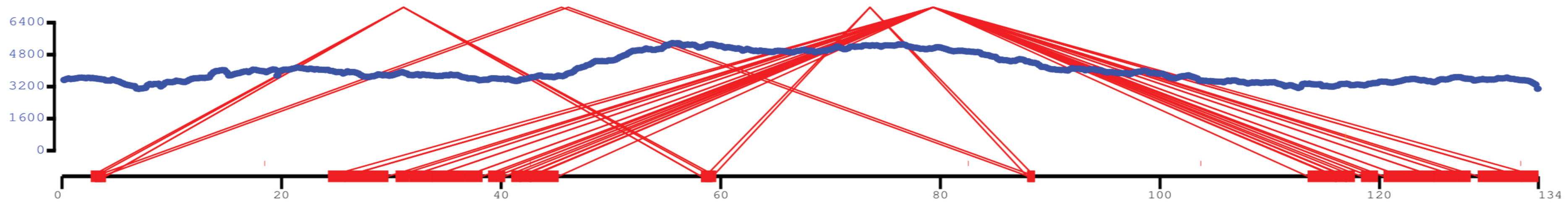

MSMg01-355O22

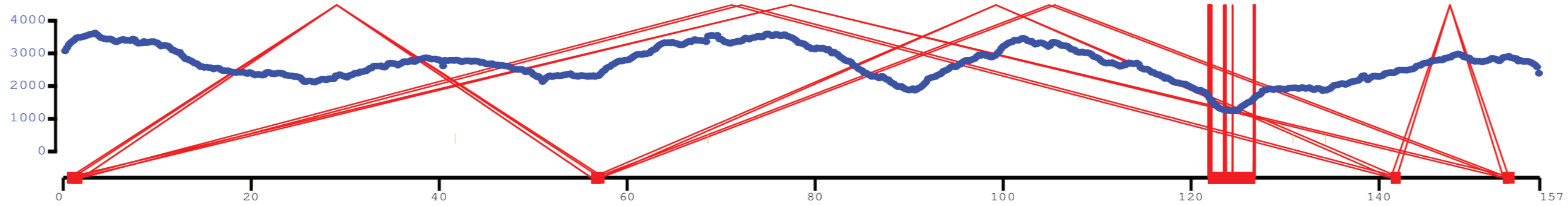


MSMg01-150D05

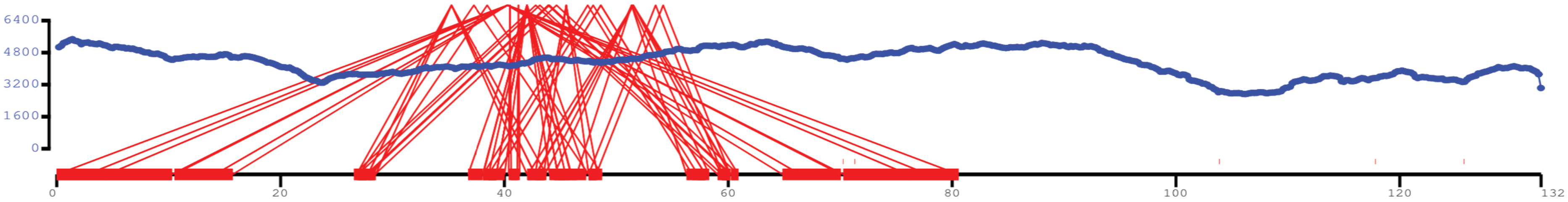


MSMg01-444A4

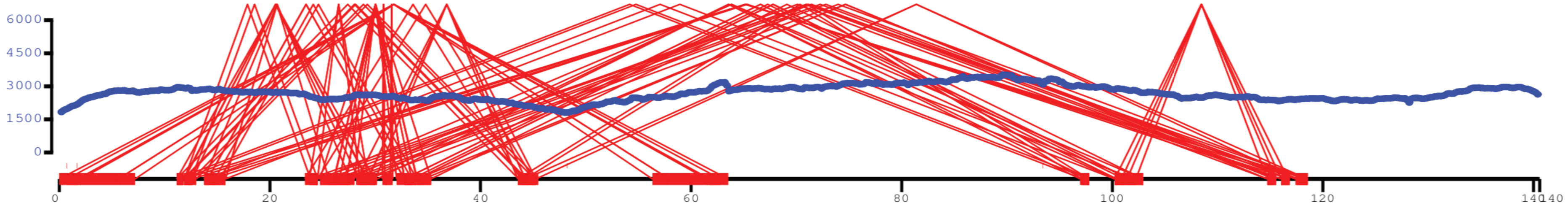


MSMg01-501C17

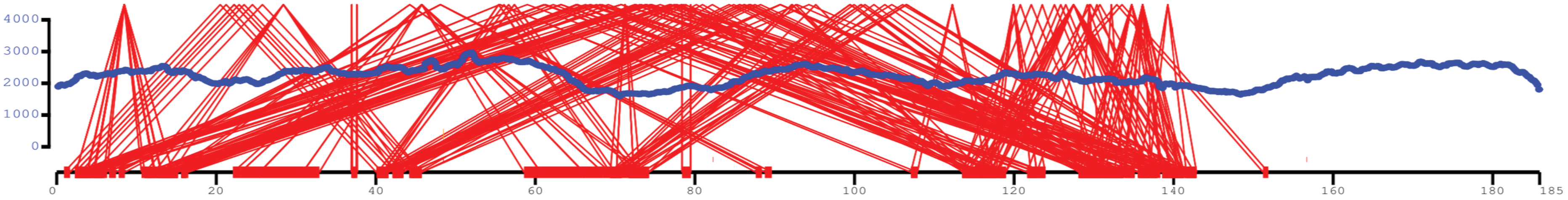

**Fig. S2**. **BAC assembly coverage visualized with Parasight**. A) Parasight images of *M. spretus* BAC assemblies. B) Parasight images of *M. molossinus* BAC assemblies. The Y axis represents fold coverage and the X axis represents length of the BAC in kb. Red lines connecting two points are internal duplications in the BAC. The blue line is coverage across the BAC assembly. Yellow tick marks above the X axis represent false alignment to the vector backbone that has been removed. Red tick marks above the X axis represent variation polished by Arrow using the raw reads. * Indicates a discontinuity in BAC CH35-113M24 due to truncated reads that fall within the palindrome arm at the end of the BAC.
